# Genetic ablation of serotonin receptor 2B improves aortic valve hemodynamics in a high-cholesterol diet mouse model

**DOI:** 10.1101/2020.08.18.255414

**Authors:** J. Ethan Joll, Cynthia R. Clark, Christine S. Peters, Michael A. Raddatz, Matthew R. Bersi, W. David Merryman

**Author notes:** **CORRESPONDING AUTHOR:** W. David Merryman, Room 9445D MRB4, 2213 Garland Ave, Nashville, TN 37212, E.

## Abstract

Calcific aortic valve disease (CAVD) is a deadly disease that is rising in prevalence due to population aging. While the disease is complex and poorly understood, one well-documented driver of valvulopathy is serotonin agonism. Both serotonin overexpression, as seen with carcinoid tumors and drug-related agonism, such as with Fenfluramine use, are linked with various diseases of the valves. Thus, the objective of this study was to determine if genetic ablation or pharmacological antagonism of the 5-HT_2B_ serotonin receptor (gene: *Htr2b*) could improve the hemodynamic and histological progression of calcific aortic valve disease. *Htr2b* mutant mice were crossed with *Notch1*^+/-^ mice, an established small animal model of CAVD, to determine if genetic ablation affects CAVD progression. To assess the effect of pharmacological inhibition on CAVD progression, *Notch1*^+/-^ mice were treated with the 5-HT_2B_ receptor antagonist SB204741. Mice were analyzed using echocardiography, histology, immunofluorescence, and real-time quantitative polymerase chain reaction. *Htr2b* mutant mice showed lower aortic valve peak velocity and mean pressure gradient – classical hemodynamic indicators of aortic valve stenosis – without concurrent left ventricle change. 5-HT_2B_ receptor antagonism, however, did not affect hemodynamic progression. Leaflet thickness, collagen density, and CAVD-associated transcriptional markers were not significantly different in any group. This study reveals that genetic ablation of *Htr2b* attenuates hemodynamic development of CAVD in the *Notch1*^+/-^ mice, but pharmacological antagonism may require high doses or long-term treatment to slow progression.

## INTRODUCTION

The aortic valve (AV) controls unidirectional oxygenated blood flow from the left ventricle into the systemic circulation. Comprised of a compliant tri-leaflet structure, this valve permits unimpeded blood flow during systole yet forms a strong coaptation to prevent regurgitation during diastole. Calcific aortic valve disease (CAVD) is characterized by the accumulation of stiff fibrotic and/or calcific deposits on the leaflet cusps which can impede blood flow, causing aortic stenosis (AS) and leading to heart failure and death if left untreated (1). AS is one of the deadliest cardiovascular diseases and is becoming an increasing public health concern in aging populations (1). The disease is estimated to be present in 1.7% of patients over the age of 65 and 3.4% over the age of 75 (2,3). Severe AS is particularly deadly, carrying a two-year survival rate as low as 22% in those with the most highly impeded blood flow (4). Estimates from 2017 indicate AS accounts for approximately 17,000 deaths in North America per year (5).

The molecular pathophysiology of CAVD is poorly understood. Originally thought to be a passive, degenerative process, research in recent decades has revealed a complex, actively regulated biological process leading to disease (6). Roles have been defined for pathological activation of resident cells including valvular endothelial cells (VECs) and valvular interstitial cells (VICs). VICs broadly contribute to CAVD through two mechanisms: dystrophic and osteogenic calcification. Dystrophic calcification is characterized by a shifting of VICs from a quiescent phenotype to a synthetic and contractile phenotype with upregulation of alpha smooth muscle actin (αSMA) and cadherin-11 (7). In osteogenic calcification, VICs can also transition to an osteoblast-like phenotype characterized by upregulation of Runt-related transcription factor 2 (Runx2), alkaline phosphatase, and ectopic bone formation (7,8).

There are currently no pharmaceutical options for slowing or treating CAVD. Surgical and transcatheter AV replacements can restore functionality; however, those suffering from late stage cardiovascular disease also often carry higher risk for complications from surgical interventions and many are left untreated (9). Therefore, a pharmaceutical alternative would be of great benefit.

One of the earliest and best-documented mechanisms driving CAVD is through the serotonin signaling pathway. Endocrine diseases leading to overexpression of serotonin, such as carcinoid tumors, have been linked with multiple valvulopathies, and serotonin receptor agonists such as Fenfluramine (an anti-obesity drug marketed in the 1990s) have been clearly shown to exacerbate valve disease (10,11). Fenfluramine was found to specifically activate 5-HT_2B_, the serotonin receptor 2B subtype (12,13). Signaling through 5-HT_2B_ has been implicated in several small animal models of cardiovascular disease. Prior work has shown that activating the 5-HT_2B_ receptor via selective agonism can lead to valve pathology in a rat model (14). Additionally, 5-HT_2B_ ablation in mice causes decreased ventricular mass and diminished systolic function (15). 5-HT_2B_ signaling has also been shown to be involved in isoproterenol and angiotensin II induced cardiac hypertrophy, primarily affecting myocardial fibroblasts (16,17). Our laboratory has previously shown that antagonism of 5-HT_2B_ can reduce AV myofibroblast activation by repressing non-canonical TGF-β signaling *in vitro* (18), indicating that 5-HT_2B_ may be a therapeutic target for CAVD.

These clinical and basic research findings led us to hypothesize that 5-HT_2B_ may be a novel target for pharmacological treatment of CAVD. Therefore, we evaluated the impact of both genetic and pharmacological ablation of 5-HT_2B_ in the well-established *Notch1*^+/-^ mouse model of CAVD, based on the heritable form in patients with *NOTCH1* mutations (19–23). Briefly, we found that genetic ablation mitigated CAVD phenotypes, while pharmacological targeting was not efficacious.

## MATERIALS AND METHODS

### Mouse models

All animal procedures were approved and carried out in accordance with relevant guidelines and regulations established by the Institutional Animal Care and Use Committee at Vanderbilt University (Protocol ID: M1600031). Briefly, *Notch1*^+/-^ and *Htr2b*^-/-^ mice were crossed to create *Notch1; Htr2b* double mutant animals (24,25). In a separate cohort, Alzet minipumps (model 1004) delivering either the high-affinity 5-HT_2B_ antagonist SB204741 (Tocris Biosciences; 1 mg kg^-1^d^-1^) or vehicle (50% dimethyl sulfoxide [Sigma-Aldrich] and polyethylene glycol-400 [Fisher Chemical]) were implanted subcutaneously in *Notch1*^+/-^ mice at 4 months of age (**Fig. 1**).

**Figure 1.**
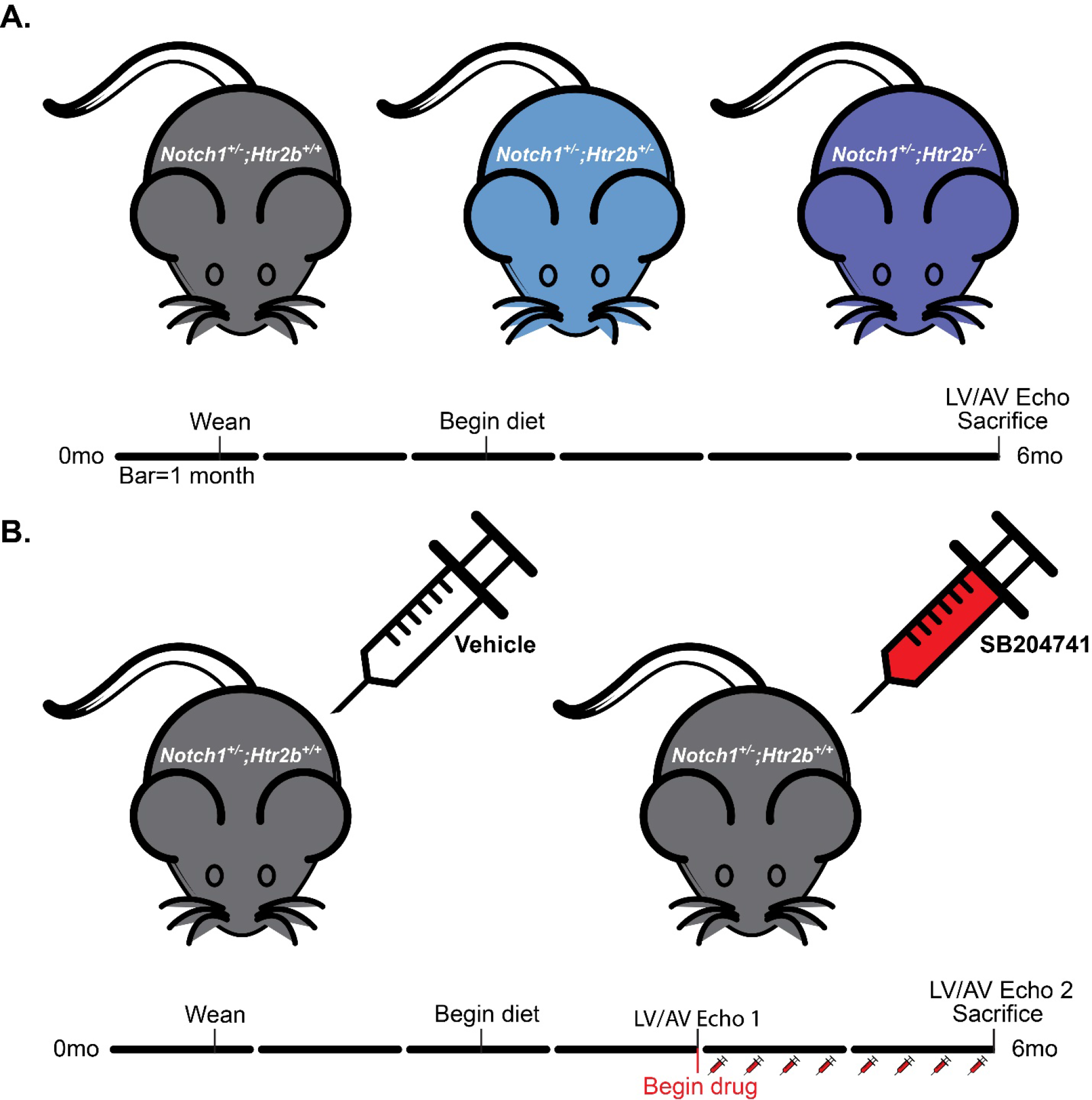
The effects of genetic ablation and pharmacological inhibition of 5-HT_2B_ on CAVD development were studied using a high-cholesterol diet *Notch1*^+/-^ mouse model. **A.** Mutant mice were aged to six months on high-cholesterol diet and doppler and M-mode echocardiography was utilized to determine hemodynamic phenotype in the valve and left ventricle. **B.** *Notch1*^+/-^ mice were aged to six months with SB204741 infusion between 4 and 6 months. Doppler and M-mode echocardiography were performed and 4 and 6 months. All mice were sacrificed at 6 months. Black bar = 1 month.

All mice were fed a 1% cholesterol Western diet (TestDiet 5TJT) for a total of 3.5 months beginning at 2.5 months of age. Food and water were provided ad libitum. Mice were sacrificed at 6 months of age using carbon dioxide inhalation followed by cervical dislocation before harvesting biological samples for processing and analysis.

### Echocardiography

*In vivo* transthoracic echocardiography was performed by the Vanderbilt Cardiovascular Physiology Core using a Vevo 2100 small animal imaging system. Mice were lightly anesthetized (average heart rate of 465 bpm) using isoflurane inhalation and placed supine on a heated stage in order to obtain a clear image of the aortic root. AV jet velocity profiles were estimated by aortic pulsed-wave Doppler imaging. A custom MATLAB script was implemented to determine the maximum systolic velocity and mean pressure gradient (26). Three images were analyzed per mouse yielding a total of ~100 cardiac cycles that were quantified and averaged to reliably quantify AV hemodynamic characteristics.

Concurrently, parasternal short-axis M-mode imaging was performed to study the effects on cardiac function and left ventricular performance. Systolic and diastolic measurements were performed to obtain estimates of ejection fraction and ventricular mass. For each mouse, ~9 cardiac cycles were averaged to obtain representative cardiac function metrics. Electrocardiographic measurements were acquired while imaging to allow for cardiac cycle-based gating of measurements using R wave peaks.

### Histological and Immunofluorescence Staining

Mice were euthanized at 6 months of age and aortic roots were immediately dissected and either flash frozen or embedded in optimal cutting temperature (OCT) compound. Embedded samples were stored at −80° C until they were cut at 10 μm serial sections in a −20° C cryostat.

To assess microstructural differences between AVs from different groups Masson’s Trichrome staining (Sigma-Aldrich) was used according to manufacturer instructions. Briefly, nuclei (black) were highlighted with Weigert’s iron hematoxylin, muscle and cytoplasm (red) with Beibrich scarlet-acid fuchsin, and collagen (blue) with aniline blue post-phosphotungstic and phosphomolybdic acid wash; slides were treated with Bouin’s solution to improve color intensity. Slides were rinsed in acetic acid, dehydrated in progressively concentrated ethyl alcohol baths, cleared with xylene, and mounted in organic mounting media. Brightfield images were taken using the Nikon Eclipse E800 microscope under a 4X magnification objective.

To assess the level of myofibroblast activation or osteoblast-like phenotype induction between groups, sections were stained for αSMA and Runx2. Sections were fixed and permeabilized in 4% paraformaldehyde/0.1% Triton-X solution. Non-specific binding was reduced by incubating in 10% bovine serum albumin in 1X phosphate buffered saline for one hour. Sections were then incubated in rabbit polyclonal anti-αSMA or rabbit monoclonal anti-Runx2 at 4° C overnight (Abcam ab5694, 1:100; Cell Signaling Technology, 1:100). Slides were thoroughly washed the next day then incubated in goat anti-rabbit IgG Alexa Fluor 647 secondary antibody for 1 hour at room temperature (ThermoFisher A-21245, 1:1000). Slides were thoroughly washed then mounted with ProLong Gold Antifade with DAPI (Cell Signaling Technology 8961), sealed, and stored at −20° C. Fluorescent images were taken using an Olympus BX53 microscope equipped with a high resolution Qimaging Retiga 3000 camera under a 20X magnification objective.

### Quantitative Image Analysis

Custom MATLAB scripts for quantitative image analysis were implemented to identify changes in leaflet composition and microstructure. Images from different samples were acquired with the same set of exposures and background levels in order to allow for quantification and comparison of image intensities.

For Masson’s Trichrome stained images, colorimetric segmentation in an HSL color space was performed in order to identify the area fractions of collagen (blue; H = 150°-250°, S = 0.1-1.0, L = 0.1-0.93) and cytoplasm/myocardium (red; H = 250°-25°, S = 0.1-1.0, L = 0.1-0.93) in each field of view (27). Area fractions for each constituent were computed as the ratio of positive pixels to non-background pixels within the specified AV leaflet area.

For immunofluorescence images, co-staining of DAPI and either αSMA or Runx2 was imaged in separate fluorescent channels at the same location. Total valve leaflet area was estimated from DAPI images based on a manually defined leaflet boundary. Segmentation of individual nuclei was also performed (28). Briefly, DAPI positive pixels were separated from the image background based on a fuzzy c-means segmentation, and individual nuclear boundaries were estimated using a modified watershed transform. Contacting nuclei were identified and split based on a concave object separation algorithm (29). αSMA area fractions were then computed as the ratio of positive pixels to total leaflet pixels based on the defined leaflet boundary. Runx2 expression was defined based on the ratio of positive pixels to total pixels within each identified nuclear boundary where Runx2 positive nuclei were defined as having greater than 50% Runx2 stain coverage. Positive nuclei counts were normalized to the number of total nuclei. αSMA and Runx2 positive pixels were defined as those with intensity values greater than the median stain intensity value within the leaflet boundary.

### Real-time Quantitative Polymerase Chain Reaction

Flash frozen aortic roots containing intact AVs were suspended in Trizol Reagent (ThermoFisher 15596026) and homogenized using a bead homogenizer (BioSpec Products). RNA was isolated using phenol-chloroform extraction as described based on manufacturer protocols. RNA purity and concentration were measured using the NanoDrop UV Spectrophotometer (ThermoFisher ND-ONE-W). cDNA was generated with the SuperScript IV First-Strand Synthesis System (ThermoFisher 18091150) based on manufacturer protocols. SYBR Green PCR Master Mix (ThermoFisher 4309155) was used to facilitate detection during target amplification. The CFX96 Real-Time PCR Detection System (Bio-Rad) was used to monitor fluorescent intensity during amplification. cDNA was initially denatured at 95° C for four minutes, then cycled 40 times through the following cycle: 10 sec 95° C, 10 sec at 55° C anneal, plate read. A melt curve was generated after cycling by increasing temperature from 65 to 95° C at 0.5° C for 5 sec and performing a plate read at each increment. Bio-Rad CFX Manager 3.1 software was used to automatically calculate the cycle quantification value at which samples amplified at a high enough value to be detected. Relative changes in transcript expression were determined by normalizing to the geometric mean of two housekeeping genes: *Gapdh* and *Actb*. Statistics were performed on untransformed ΔC_t_ values. All primers are included below.

Primers were designed using the Integrated DNA Technologies PrimerQuest tool. Primers were designed to span one exon and with a 3’ bias. Product sizes of less than 300 were selected. Sequences were verified using NCBI Primer-BLAST to fully match the target transcript and have no significant off target interactions.

**Table 1.**
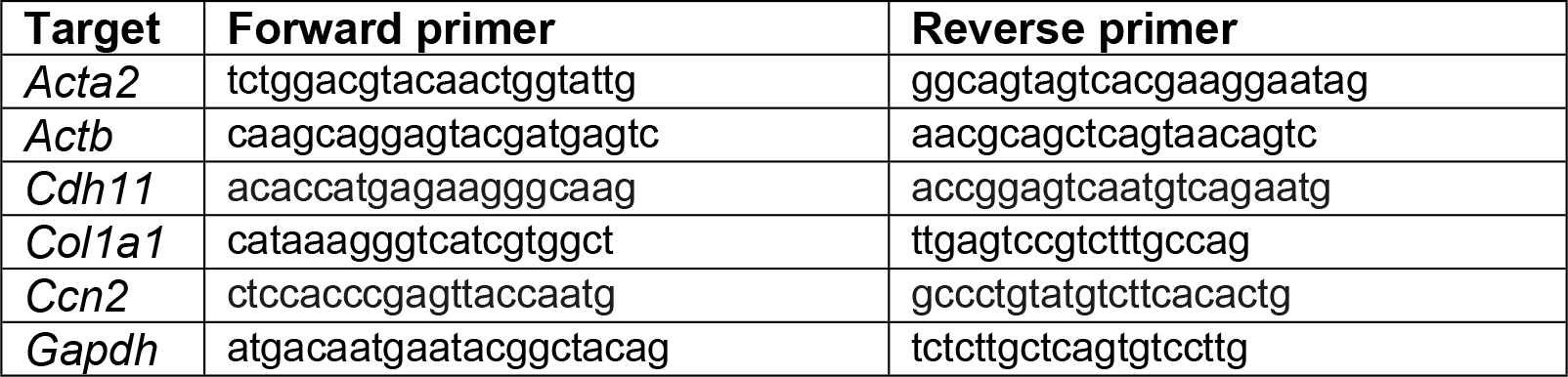
Primer nucleotide sequences used for RT-qPCR.

### Statistical Analysis

Data are presented as mean +/- standard error of the mean (SEM), unless otherwise noted. All data sets were tested for normality using the D’Agostino-Pearson omnibus normality test. Data sets passing the normality test were analyzed using parametric tests: Student’s *t*-test for 2 group comparison and one-way analysis of variance (ANOVA) for 3 or more groups. Small groups (sample size less than 7) or groups with non-normal distribution were analyzed using non-parametric tests: Mann-Whitney U test for 2 group comparison and Kruskal-Wallis one-way analysis of variance for 3 or more groups. For analyzing absolute change from four to six months in the drug study, the one-sample *t* test or Wilcoxon signed-rank test was used for parametric and non-parametric data, respectively. To account for multiple comparisons, statistical significance was corrected in post-hoc tests using Tukey or Dunn’s correction for parametric and non-parametric tests, respectively. For all tests, a p-value of 0.05 was considered statistically significant. Data storage and statistical analysis was performed using Microsoft Excel, MATLAB r2019a, and GraphPad Prism 8.4.1.

## RESULTS

### Genetic ablation of Htr2b rescues hemodynamic metrics of aortic valve disease

Ablation of *Htr2b* from *Notch1*^+/-^ mice decreases the average peak velocity during the cardiac cycle with a corresponding decrease in mean pressure gradient across the AV, relative to *Notch1*^+/-^;*Htr2b*^+/+^ (**Fig. 2a-b**). Left ventricular mass was not changed with *Htr2b* ablation. (**Fig. 2c**). Additionally, the function of the left ventricle was not altered based on ejection fraction measurements (**Fig. 2d**). Together, this suggests the observed changes in hemodynamics are intrinsic to the AV and not due to altered cardiac function.

**Figure 2.**
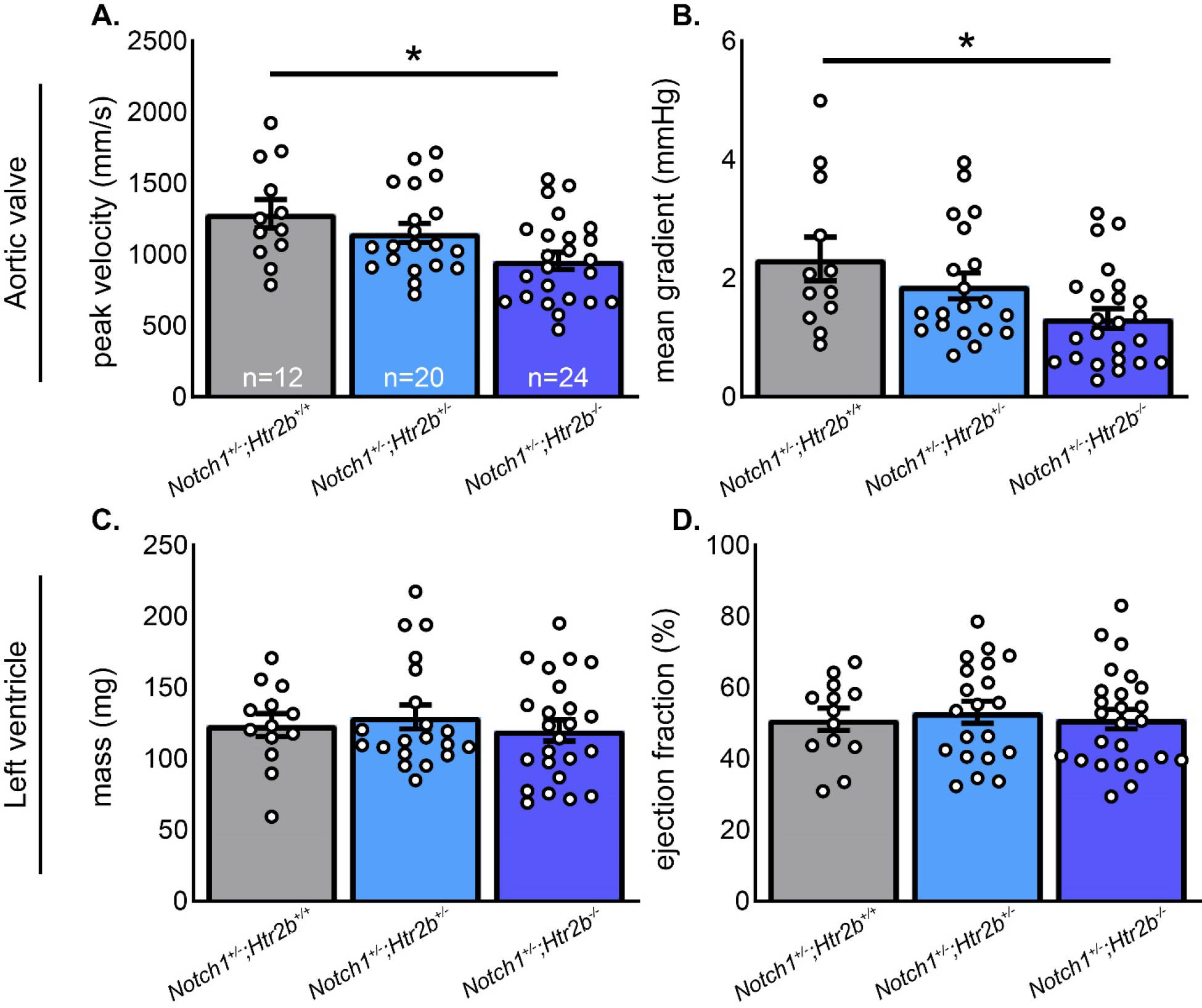
Genetic ablation of 5-HT_2B_ results in a valve-specific hemodynamic improvement. Sequential reduction of *Htr2b* expression from *Notch1*^+/-^ mice results in a corresponding reduction in **A.** peak systolic velocity (mm/s) and **B.** mean pressure gradient (mmHg) across the AV at 6 months of age. Left ventricular structure and function were assessed by measurements of **C.** mass (mg) and **D.** ejection fraction (%), respectively. Neither metric was altered in *Htr2b* mutants indicating a primarily valvular phenotype. Sample sizes for all groups labeled in white in panel **A.** Mean+/-SE, *p<0.05, 1-way ANOVA with post-hoc Tukey test for multiple comparisons.

### 5-HT_2B_ antagonism does not affect hemodynamic metrics of aortic valve disease

Based on results from the mutant study, we aimed to determine if the beneficial effects of *Htr2b* ablation on CAVD progression could be recapitulated using a pharmacological inhibitor specific to the 5-HT_2B_ receptor. Following the initiation of SB204741 treatment (after 6 weeks of high cholesterol diet), the absolute change in peak velocity, mean gradient, left ventricular mass, and ejection fraction was calculated from 4 to 6 months of age. There was no significant change in valve metrics of peak velocity or mean pressure gradient **(Fig. 3a-b)**. Further, there was a slight increase in left ventricle mass but no significant change in ejection fraction **(Fig. 3cd)**. Despite AV improvement following genetic ablation, 5-HT_2B_ antagonism by SB204741 does not reduce CAVD progression in *Notch1*^+/-^ mice.

**Figure 3.**
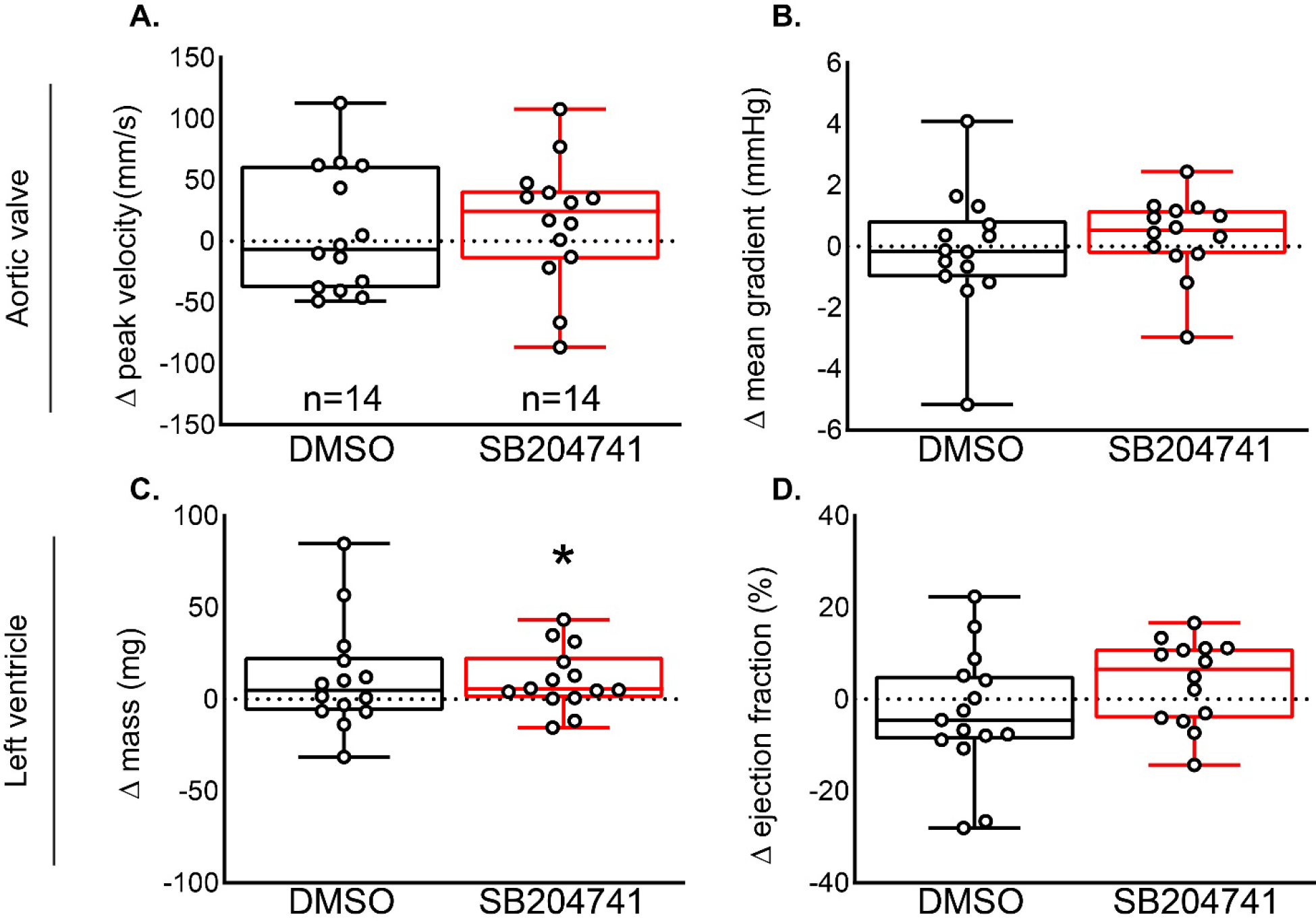
Pharmacological inhibition of 5-HT_2B_ by SB204741 does not improve AV hemodynamics between four and six months. Change in **A.** peak velocity and **B.** mean gradient revealed no differences in AV hemodynamic metrics between four and six months for DMSO or SB204741 treated groups. **C.** Left ventricle mass increased slightly in drug treated mice, however this did not translate to a statistically significant change in **D.** ejection fraction between four and six months. Boxes extend from mean to 25 and 75 percentiles and whiskers to minimum and maximum values, *P<0.05, (**A, D**) one sample T-test or (**B, C**) Wilcoxon signed-rank test.

### 5-HT_2B_ genetic ablation or pharmacological inhibition does not significantly change leaflet microstructure

Masson’s trichrome staining was used to analyze collagen abundance and thickness of AV leaflet cross sections. There was wide variation across samples and despite trends toward decreased thickness and increased collagen density, we observed no statistically significant changes (**Fig. 4a**). Similarly, there were no detected differences in leaflet thickness or collagen density in drug treatment groups despite the same trends as observed in the genetic ablation study (**Fig. 4b**).

**Figure 4.**
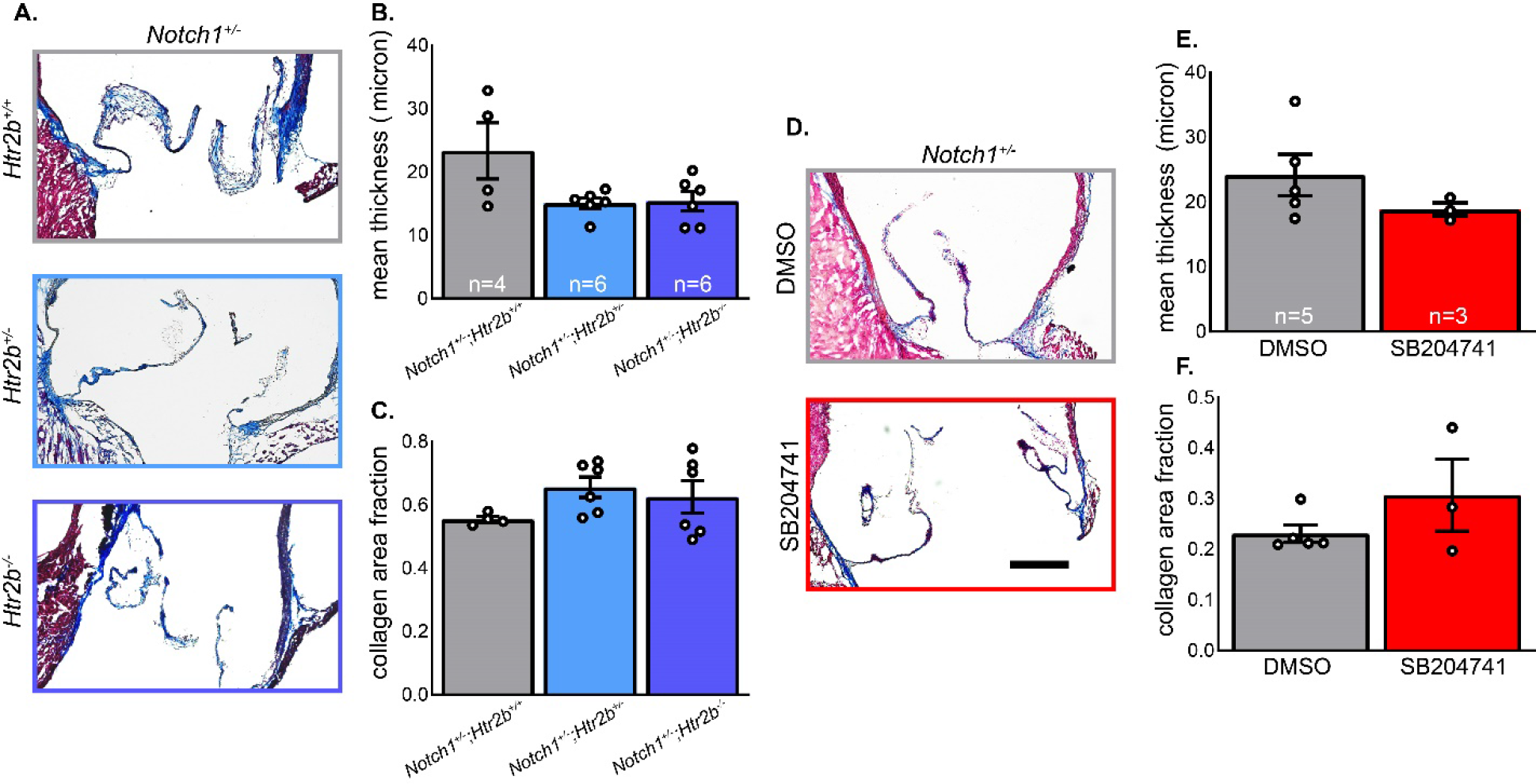
Neither 5-HT_2B_ genetic ablation nor pharmacological inhibition induces changes in leaflet thickness or collagen density. **A.** Masson’s trichrome stain was used to distinguish valve area and collagen tissue in mutant mice. **B.** Mean leaflet thickness was determined by measuring total valve area normalized length along cusp and was not changed. **C.** Collagen area fraction was not significantly changed between groups. **D.** Masson’s trichrome stain was used in drug and vehicle treated mice. **E.** Mean leaflet thickness was not changed between treatments. **F.** Collagen density was not significantly different between treatments. Mean+/-SE, *P<0.05, (**B, C**) Kruskal-Wallis 1-way ANOVA with post-hoc Dunn’s test, (**E, F**) Mann-Whitney U-test. Black bar = 200 μm.

### 5-HT_2B_ genetic ablation or pharmacological inhibition does not significantly change myofibroblast activation or osteoblast-like phenotype induction

To determine if myofibroblast activation played a role in the difference observed in mutant AV hemodynamic phenotypes, immunofluorescent labeling of the αSMA protein was performed (**Fig. 5a-b**). Similarly, Runx2 was targeted to determine if there was evidence of osteoblast-like phenotype shift in valve cells (**Fig. 6a-b**). There was no significant difference in αSMA and high sample-to-sample variance was observed, particularly in the *Notch1*^+/-^; *Htr2b*^+/+^ group (**Fig. 5c**). Analysis of Runx2 staining revealed a 50-60% positivity rate of all valve cells independent of genotype and treatment. No significant differences in Runx2 expression were observed between groups (**Fig. 6c**).

**Figure 5.**
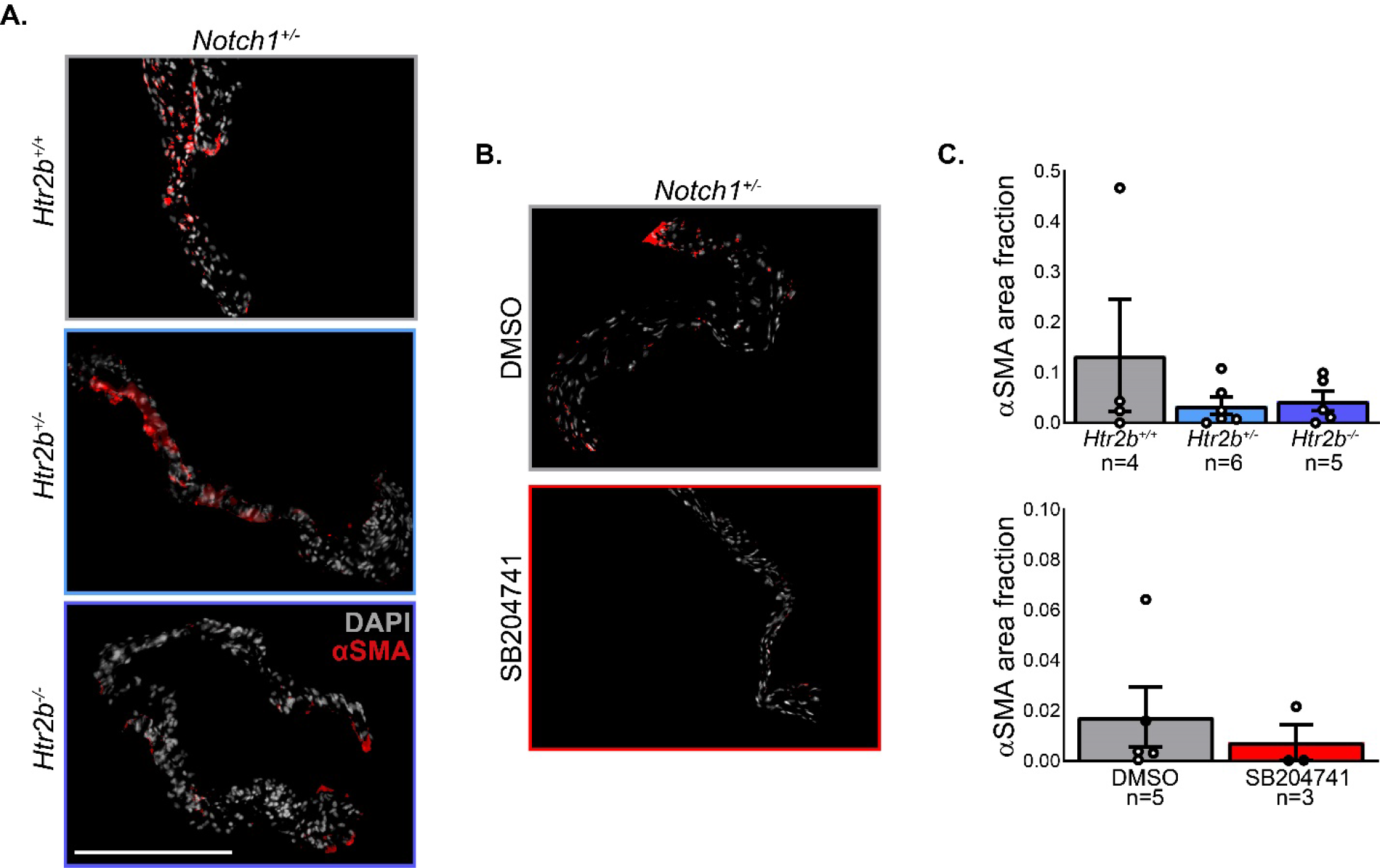
Neither 5-HT_2B_ genetic ablation nor pharmacological inhibition show a difference in myofibroblast activation based on αSMA positive valve area. **A.** Fluorescent imaging was carried out on mutant mice in the far red and DAPI channels for αSMA and nuclei imaging, respectively. **B.** Fluorescent imaging was carried out as described on vehicle and drug treated mice as described. **C.** Area positive for αSMA stain was normalized to the total valve area calculated by DAPI masking. Mean+/-SEM, *p<0.05 following Kruskal-Wallis 1-way ANOVA with Dunn’s post-test. White bar = 200 μm.

**Figure 6.**
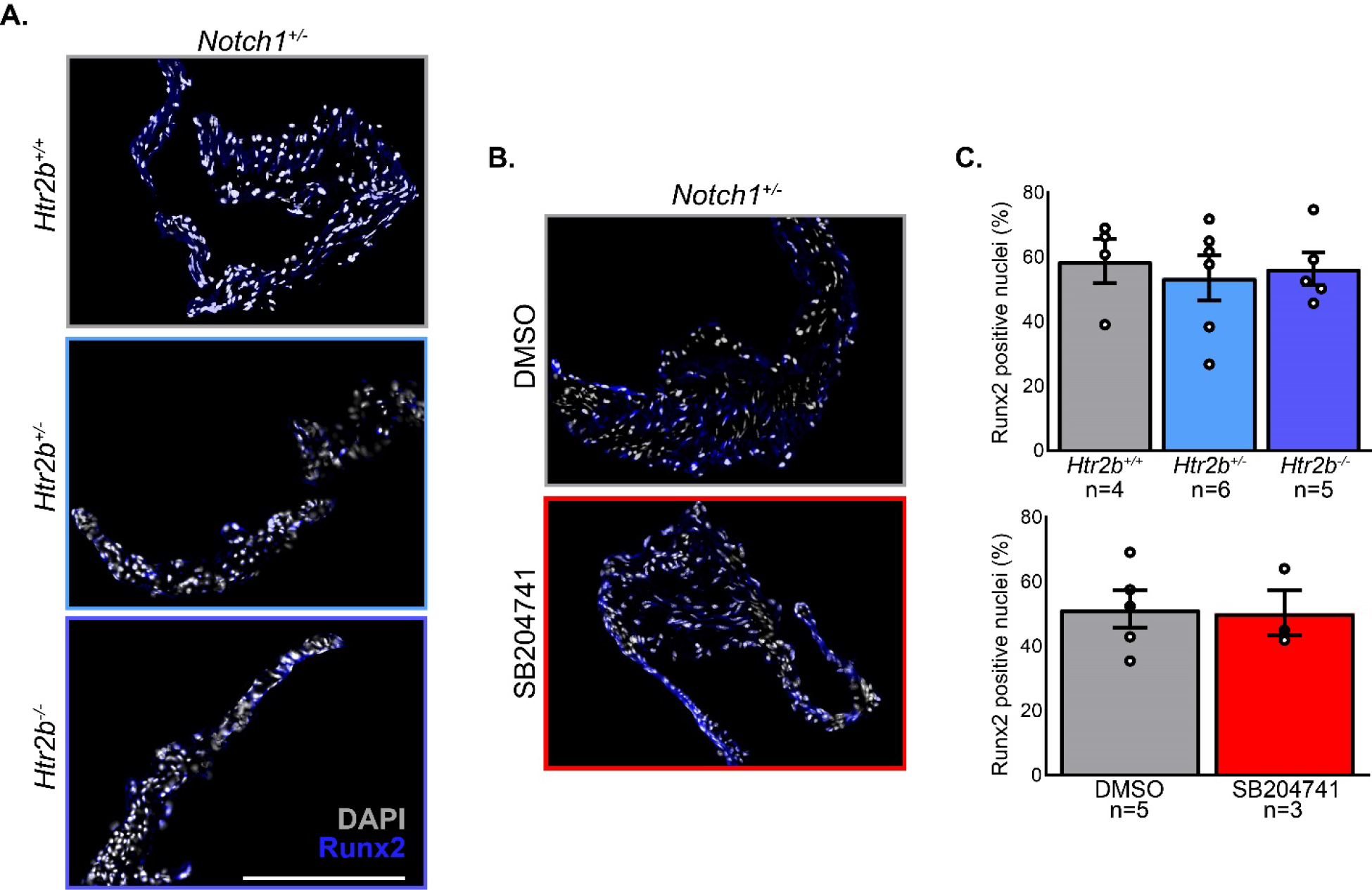
Neither 5-HT_2B_ genetic ablation nor pharmacological inhibition show a difference in osteoblast-like phenotype based on Runx2 positive nuclei. **A.** Fluorescent imaging was carried out on mutant mice in the far red and DAPI channels for Runx2 and nuclei imaging, respectively. **B.** Fluorescent imaging was carried out as described on vehicle and drug treated mice as described. **C.** Nuclei greater than 50% positive for Runx2 stain were counted and normalized by total nuclei. Mean+/-SEM, *p<0.05 following Kruskal-Wallis 1-way ANOVA with Dunn’s post-test. White bar = 200 μm.

### 5-HT_2B_ genetic ablation or pharmacological inhibition does not significantly change fibrotic CAVD transcripts

To further investigate the molecular mechanisms leading to differing hemodynamic phenotypes in mutant mice, RT-qPCR was performed for several genes of interest. Levels of these transcripts were not significantly changed, further pointing to a myofibroblast independent mechanism (**Fig. 7a**). *Ccn2*, the gene for the fibrotic marker connective tissue growth factor (CTGF) showed an overall group difference (p < 0.05; Kruskal-Wallis 1-way ANOVA) but no differences when accounting for multiple comparisons (p = 0.0655 between *Notch1*^+/-^; *Htr2b*^+/+^ and *Notch1*^+/-^; *Htr2b*^-/-^; p = 0.0826 between *Notch1*^+/-^; *Htr2b*^+/-^ and *Notch1*^+/-^; *Htr2b*^-/-^).

**Figure 7.**
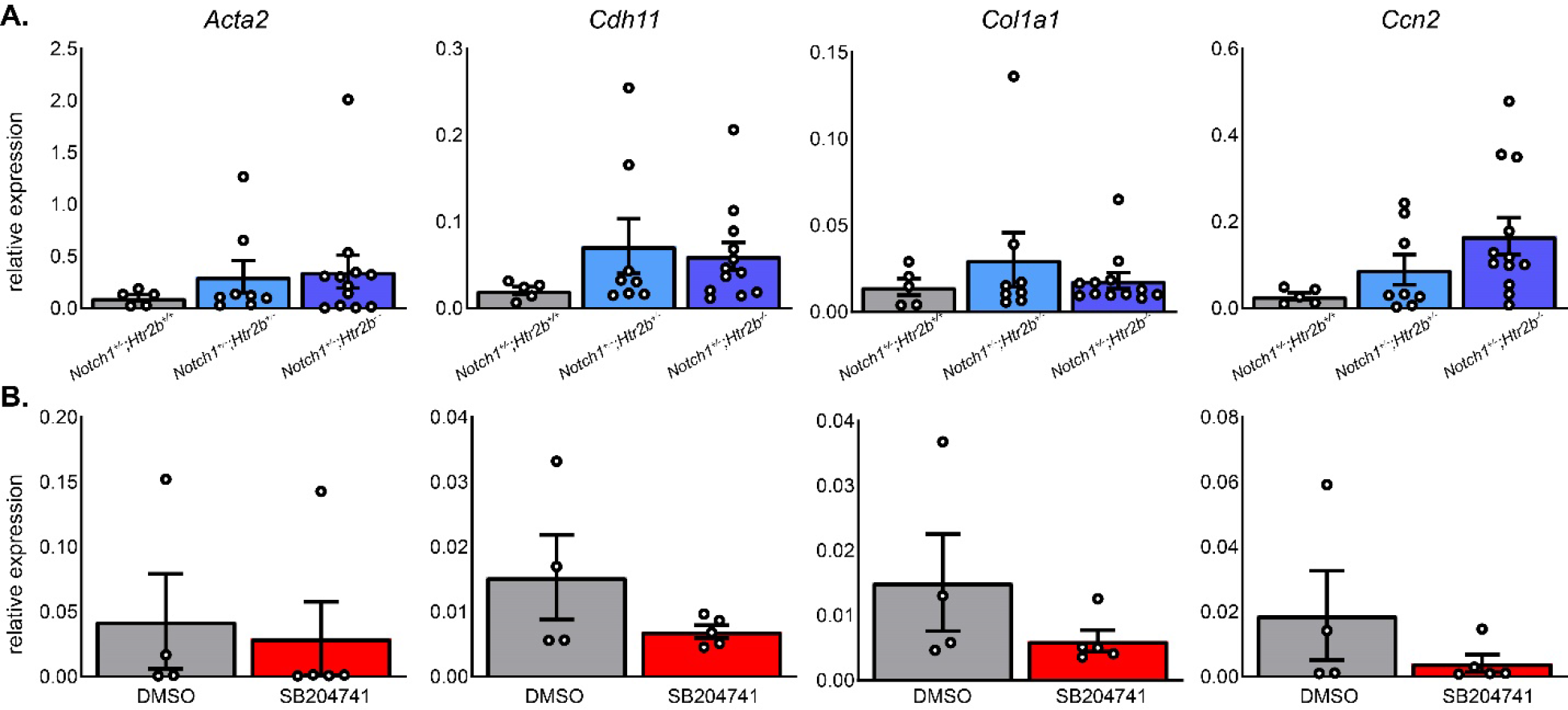
*Htr2b* mutant and SB204741-treated mice do not have altered transcription of genes associated with fibrotic/myofibroblast driven CAVD. **A.** RT-qPCR was used to assess the levels of mRNA transcripts of *Acta2, Cdh11, Col1a1*, and *Ccn2*. No significant differences were detected. **B.** Similar targets were interrogated in drug treatment groups. Similarly, there were no differences. Mean+/-SE, *P<0.05 following Kruskal-Wallis 1-way ANOVA with Dunn’s post-test.

RT-qPCR was performed in order to potentially detect any early markers of disease that may be present before symptoms can be detected by imaging. No significant changes were detected in the typical fibrotic CAVD markers of *Acta2, Cdh11, Col1a1*, or *Ccn2* (**Fig. 7b**).

## DISCUSSION

CAVD is a significant health concern in aging populations of developed countries, which are shifting toward more sedentary lifestyles. While the pathophysiology of the disease is poorly understood, serotonin signaling is a relatively well characterized driver of valve disease. The aim of this study was to assess the possibility of slowing or treating CAVD by genetically deleting or pharmacologically inhibiting 5-HT_2B_ in a high-cholesterol diet *Notch1*^+/-^ mouse model. We found that genetic mutation of the 5-HT_2B_ receptor – via *Htr2b* deletion – mitigates hemodynamic progression of aortic stenosis.

We began by testing echocardiographic outcomes of genetic and pharmacological blockade of 5-HT_2B_. Genetic knockdown of *Htr2b* improved echocardiographic metrics of aortic stenosis (**Fig. 2**), but receptor blockade with SB204741 had no effect (**Fig. 3**). This may indicate that pharmacological inhibition of 5-HT_2B_ using SB204741 is insufficient in blocking receptor action compared to complete genetic deletion. This is supported by the fact that *Notch1*^+/-^; *Htr2b*^+/-^ mice saw only slight, non-significant improvement over *Notch1*^+/-^; *Htr2b*^+/+^ mice. In other words, only complete *Htr2b* knockout mice had significant changes. A stronger hemodynamic phenotype tends to develop between 6 and 9 months of age when using HFD (unpublished observations). It is possible that the mice in our study were not aged long enough to see appreciable differences in CAVD phenotypes. The age-range in this study is comparable to early adulthood in humans whereas CAVD generally does not become prevalent until advanced age.

On assaying leaflet morphology and collagen structure, we saw no significant changes in any group (**Fig. 4**). This was expected in drug treatment studies as we saw no significant differences in hemodynamic profile; therefore, we expected the structure was likely unchanged. Trends in mutant mice may indicate reduced leaflet hypertrophy in mutants which could lead to improvements in the AV hemodynamic profile. However, variability in the studied groups – particularly *Notch1*^+/-^;*Htr2b*^+/+^ – precludes a clear conclusion from being drawn. The lack of change in collagen density may indicate an alternate structural cause for differences in the systolic AV flow profile in mutant mice, such as accumulation of glycosaminoglycans or an osteogenic calcification phenotype, undetected by using the trichrome staining method (6,8,30).

We also assessed myofibroblast activation by αSMA staining. Serotonin signaling through the 2B receptor has shown to play a role in the activation of quiescent VICs into activated myofibroblasts. These cells play a role in the fibrosis and calcification that lead to impaired blood flow found in CAVD. One of the integral parts of myofibroblast activation is the adoption of a contractile phenotype that is characterized by abundant stress fibers primarily composed of the protein αSMA (7). Yet again, we saw no significant changes (**Fig. 5**). It is possible that the myofibroblast activation phenotype may be a bimodal distribution and if a longer time course was studied, more mice would develop the elevated αSMA phenotype seen in one *Notch1^+/-^;Htr2b^+/+^* mouse. However, when combined with collagen deposition based on trichrome staining, it appears the myofibroblast-driven fibrotic phenotype is likely not the cause of the differing phenotype which may point to a difference in glycosaminoglycan structure, osteogenic calcification, or cellular inflammation (6,30,31). *Notch1* is known to downregulate Runx2 mediated osteoblast-like phenotype in CAVD (19,32). Indeed, immunostaining in this study shows Runx2 expression is present in *Notch1*^+/-^ mice with around 50% of resident cells positive for the protein. Our data show *Htr2b* ablation nor receptor inhibition alter Runx2 expression, suggesting the hemodynamic phenotype observed in mutant mice is not based on canonical osteogenic signaling pathways (**Fig. 6**). Immune cell recruitment plays a key role in other 5-HT_2B_-mediated fibrotic disease and has been shown to contribute to the *Notch1*^+/-^ model of CAVD, suggesting it may be involved here (23,33,34). This may also indicate a phenotype arising from cells other than myofibroblasts. For instance, valvular endothelial cells are involved in CAVD progression via nitric oxide signaling and endothelial-mesenchymal transition (35,36). These alternative cellular mechanisms have not been rigorously interrogated in the context of 5-HT_2B_ associated CAVD and may be a fruitful avenue of investigation.

RT-qPCR studies (**Fig. 6**) also reveal little about the causative mechanisms of the changes found in the hemodynamics of the mutants. Changes in myofibroblast markers *Acta2* and *Cdh11* were not changed in mutants or drug groups. *Col1a1* was also not altered in any group, as would be expected based on histology analysis. Finally, *Ccn2*, a marker of fibrotic remodeling, was different at a group level in mutants but after correcting for multiple comparisons we observed no differences. This was also true between drug treatment groups.

Overall, our results indicate that double mutant mice exhibit an improved hemodynamic phenotype at six months of age compared to *Notch1*^+/-^; *Htr2b*^+/+^ mice. However, pharmacologic inhibition of 5-HT_2B_ by SB204741 did not affect the development of CAVD between four and six months of age. This may be due to either antagonism of the receptor not being a sufficient intervention to slow progression of the disease or our window of intervention may have been too early as six months of age is still relatively early for the disease to present. Indeed, results from the genetic study suggest prophylactic 5-HT_2B_ inhibition may be sufficient for attenuating CAVD initiation, but therapeutic inhibition is insufficient for slowing the progression of existing disease. Histological analysis revealed mutant and drug treated mice did not significantly vary in level of leaflet hypertrophy or collagen density. There was also no difference in a subset of markers associated with myofibroblast activation and fibrosis that are generally detected in CAVD. Indeed, at the transcript level, there was little difference in general markers of CAVD.

Findings from the current study highlight the involvement of 5-HT_2B_ signaling on CAVD initiation and progression based on improved AV hemodynamics following *Htr2b* deletion. Future studies focused on the therapeutic potential of targeted 5-HT_2B_ receptor blockers should include a longer time course (12-18 months) to study the impact on the later stages of CAVD where the hemodynamic, tissue, and molecular phenotypes should be stronger and interrogate the effect of 5-HT_2B_ inhibition before and after disease onset. Additionally, unbiased transcriptomic and proteomic approaches, such as mRNA sequencing, can be used to identify potential pathways of interrogation that may have been missed using a targeted approach.

### Limitations

We were not able to age mice past 6 months in this study, limiting the development of CAVD and AS phenotypes which correlate strongly with advanced age. Molecular interrogation of causative pathways was limited due to the small amount of high-quality mRNA and tissue sections that can be extracted from single mouse aortic roots.

## CONCLUSIONS

The results from this study provide evidence that *Htr2b* mutant mice may have more compliant leaflets allowing for unimpeded blood flow in the *Notch1*^+/-^ model of CAVD. However, the molecular changes underlying this tissue-level difference are unclear. We also found that despite significant improvement following genetic deletion, 5-HT_2B_ antagonism using SB204741 was unable to cause a significant hemodynamic or histological difference between four and six months of age. Future studies are needed to fully evaluate the therapeutic potential of targeting 5-HT_2B_ signaling for the treatment and management of CAVD.

## ACKNOWLEDGEMENTS

This work was supported by the following NIH grants: F31-HL151115 (JEJ II), F30-HL147464 (MAR), K99-HL146951 (MRB), R01-HL115103 (WDM), R35-HL135790 (WDM). The funders had no role in study design, data collection and analysis, decision to publish, or preparation of the manuscript.

